# IAPP Marks Mono-hormonal Stem-cell Derived β Cells that Maintain Stable Insulin Production *in vitro* and *in vivo*

**DOI:** 10.1101/2024.04.10.587726

**Authors:** Jeffrey C. Davis, Maria Ryaboshapkina, Jennifer H. Kenty, Pınar Ö. Eser, Suraj Menon, Björn Tyrberg, Douglas A. Melton

## Abstract

Islet transplantation for treatment of diabetes is limited by availability of donor islets and requirements for immunosuppression. Stem cell-derived islets might circumvent these issues. SC-islets effectively control glucose metabolism post transplantation, but do not yet achieve full function *in vitro* with current published differentiation protocols. We aimed to identify markers of mature subpopulations of SC-β cells by studying transcriptional changes associated with *in vivo* maturation of SC-β cells using RNA-seq and co-expression network analysis. The β cell-specific hormone islet amyloid polypeptide (IAPP) emerged as the top candidate to be such a marker. IAPP^+^ cells had more mature β cell gene expression and higher cellular insulin content than IAPP^-^ cells *in vitro*. IAPP^+^ INS^+^ cells were more stable in long-term culture than IAPP^-^ INS^+^ cells and retained insulin expression after transplantation into mice. Finally, we conducted a small molecule screen to identify compounds that enhance IAPP expression. Aconitine up-regulated IAPP and could help to optimize differentiation protocols.

**Highlights:**

- IAPP expression *in vitro* marks a mono-hormonal subpopulation of SC-β cells excluding endocrine hormones other than insulin
- Only INS^+^ IAPP^+^ cells maintain stable *INS* expression *in vitro* up to 100 days after differentiation
- The small molecule aconitine accelerates IAPP expression in SC-β cells *in vitro*

## Introduction

Diabetes is a heterogenous disease affecting 451 million people worldwide (Cho et al., 2018). Pancreatic β cell death and dysfunction are the primary pathophysiological mechanisms in patients with severe autoimmune diabetes and severe insulin-deficient diabetes, comprising 6.4% and 17.5% of the diabetic population, respectively (Ahlqvist et al., 2018). β cell transplantation therapies have the potential to replace insulin injections and offer a cure for diabetic patients. However, islet transplantation is restricted by availability of high-quality donor islets and the requirement for immunosuppression to achieve graft survival (Krentz et al., 2021). Human stem cell-derived β (SC-β) cells are an emerging alternative cell source for transplantation that could overcome both of these obstacles (Krentz *et al*., 2021). SC-β cells secrete insulin in response to glucose and rescue glycemic control after transplantation into diabetic rodents (Pagliuca et al., 2014; Rezania et al., 2014; Russ et al., 2015; Velazco-Cruz et al., 2019).

SC-β cells are not yet fully functional at the end of *in vitro* differentiation using currently published protocols to generate SC-islets from human pluripotent stem cells (Davis et al., 2020). However, SC-β cells mature when transplanted under the kidney capsule of immunocompromised mice (Pagliuca *et al*., 2014). Currently, SC-β cells for preclinical transplantation studies are defined by co-expression of the β cell transcription factor NKX6-1 and the C-peptide fragment of processed insulin (Pagliuca et al., 2014; Veres et al., 2019). These markers exclude poly-hormonal cells with inappropriate non-β cell hormone expression and select cells that remain insulin-positive after transplantation. NKX6-1 is traditionally considered an essential and specific β cell marker (Schaffer et al., 2013). However, recent single-cell RNA-seq data indicate that NKX6-1 is also expressed by progenitor (immature) cells and enterochromaffin cells generated during *in vitro* differentiation (Veres *et al*., 2019). Alternative β cell enrichment strategies are based on staining for intracellular zinc (Davis et al., 2019; Jindal et al., 1993; Lukowiak et al., 2001) and ITGA1 (Veres *et al*., 2019). However, ITGA1 is not specific to adult functionally mature human β cells in islets (Baron et al., 2016). Zinc granules may also be contained in immature fetal β cells (Meyramova et al., 2021; Pound et al., 2012). Thus, more specific markers are needed to identify functionally mature β cells among the SC-β population. This study aims to identify such markers.

## Results

### Identification of IAPP

We first explored how changes in gene expression might correlate with improved glucose-stimulated insulin secretion (GSIS) after engraftment. The 1016 iPS^GFP^ cell line constitutively expresses GFP from the CAG promoter, enabling the isolation of iPS derived SC-islets following transplantation into mice. 1016 iPS^GFP^ cells have a modest efficiency in generating NKX6-1^+^ C-peptide^+^ cells when differentiated *in vitro* to SC-islets using a 6-stage protocol (Pagliuca *et al*., 2014). We isolated the SC-islets using GFP expression, the live cell zinc dye TSQ, and negative selection for propidium iodide uptake (Davis *et al*., 2019), in the same differentiations before and 6 weeks after transplantation **(Figure 1A).** Post-transplantation SC-β cells lost their poly-hormonal cell population, resolving into mono-hormonal α- and β-like cells (**Figure 1B**). Basal insulin secretion during fasting remained unchanged (**Figure 1C**) while GSIS improved following *in vivo* maturation (**Figure 1D**). Levels of most islet endocrine hormones remained unchanged. The β cell hormones, islet amyloid polypeptide (IAPP), and adenylate cyclase activating peptide (ADCYAP1) were up-regulated after transplantation (**Figure 1E**). Co-expression network analysis in human islets revealed that IAPP and ADCYAP1 belonged to the same module (**Figure 1F**). This gene community was enriched in differentially expressed genes after transplantation (p 2e-7) and genes with calcium ion binding activity (GO:0005509, p 3e-6). The first principal component (PC1) correlated with other studies that investigated islet gene expression signatures associated with HbA1c (Kendall tau 0.29 p 2.7e-4) and BMI (tau 0.22 p 2.9e-3) but not age (tau −0.09, non-significant) supporting functional importance. IAPP had the highest contribution to the first principal component (**Figure 1F**) and was selected for further experiments.

**Figure 1.**
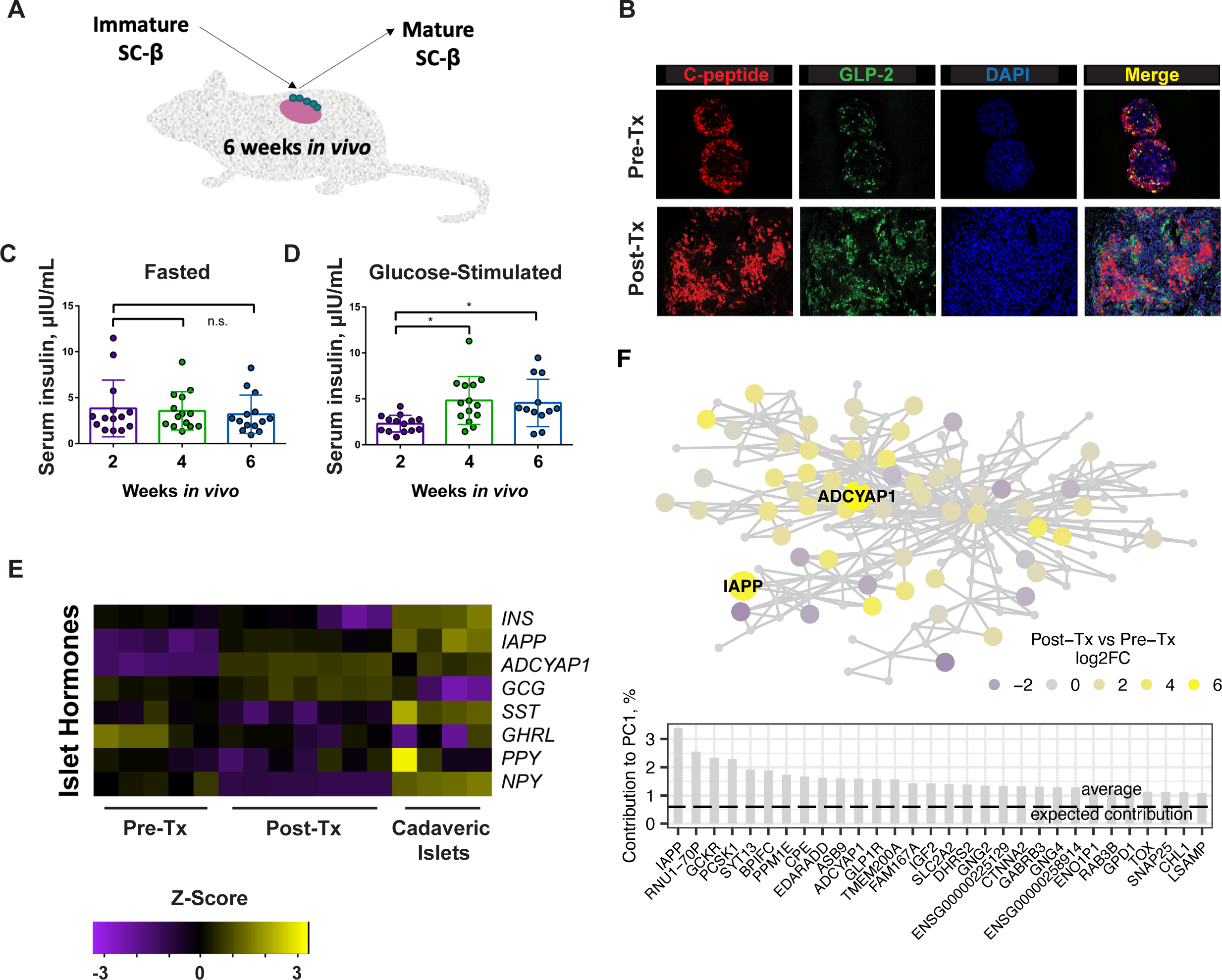
Identification of IAPP. (**A)** Outline of the *in vivo* maturation experiment transplanting SC-islets to the kidney capsule in immunodeficient mice. (**B**) Immunofluorescence of β cell hormone C-peptide and α cell GLP-2 before and after transplantation (Tx) of SC-islets indicates lack of polyhormonal cells in grafts. Human insulin levels in (**C**) fasted and (**D**) glucose-stimulated state at three timepoints during *in vivo* engraftment. Statistical significance was determined using a 2-way ANOVA test (p < 0.01.) (**E**) Islet hormone expression in SC-islets before and after transplantation by RNA-seq. n = 5 differentiations pre-transplant, n = 7 mice post-transplant, and n = 4 cadaveric islet donors. (**F**) Network-based prioritization of IAPP. The schematic graph shows gene community (circles) and its underlying correlation structure (lines connecting the dots). Circle color indicates fold change after vs before transplantation. Circles signed with gene symbol are β-cell enriched genes. Bar graph illustrates top 30 member genes by contribution to the first principal component (PC1).

### Dual INS and IAPP Reporter Line

We generated a dual INS and IAPP knock-in reporter line in the 1016 iPS cell background. An Insulin Red Nucleus (IRN) INS^mCherry^ knock-in line was generated by inserting an in-frame mCherry coding sequence downstream of the *INS* gene, separated by a P2A cleavage sequence (**Figure 2A**). Specificity was confirmed using real time RT-PCR as well as genomic PCR. A single clone of IRN 1016 iPS cells was chosen after validation and further targeted with a knock-in construct to generate an IAPP^eGFP^ knock-in line, Amylin Nuclear Green in IRN (ANGIs) (**Figure 2B**). Thus, this iPS cell line, ANGIs, expresses mCherry in the cytoplasm when the insulin gene is expressed and, amylin nuclear green when the IAPP gene is expressed. A fraction of the INS^+^ population expressed IAPP on day 7 of the final, 6^th^ stage of differentiation *in vitro* (**Figure 2C**). IAPP expression was nearly absent outside of the INS^+^ population (**Figure 2C**). Epifluorescence analysis of live SC-islets revealed distinct localization of the IAPP^eGFP^ in the nucleus and INS^mCherry^ in secretory granules (**Figure 2D**). While mCherry localization was not nuclear as expected, the reporter line was still able to mark insulin-expressing cells and secreted insulin was detectable by ELISA in subsequent experiments. The final stage of SC-islet differentiation media is composed of basal media without additional growth factors following successful endocrine induction. INS expression peaked after 2 weeks of factor withdrawal and decreased throughout the next 3 weeks (**Figure 2E**). Yet, the proportion of IAPP^+^ cells among the remaining INS^+^ cells gradually increased with time in culture and continued even 50 and 100 days after growth factor withdrawal, as the few cells to remain INS^+^ with extended *in vitro* culture expressed IAPP (**Figure 2F**). This suggests that INS^+^ IAPP^+^ cells in stage 6 are stable over long periods in culture and account for the majority of cells that remain INS^+^ after growth factor withdrawal.

**Figure 2.**
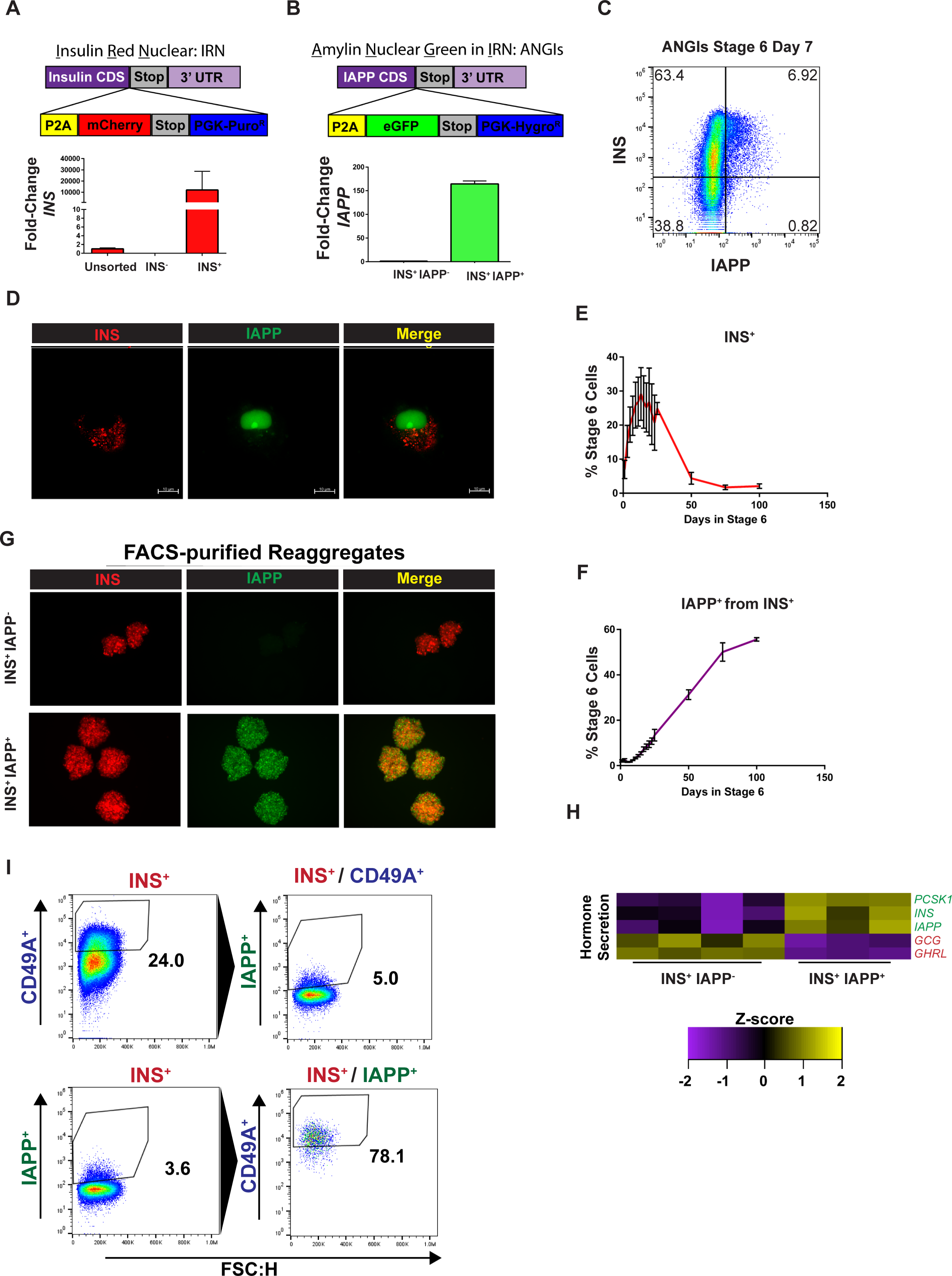
Characterization of the Dual INS and IAPP Knock-in Reporter Line ANGIs. Transgene design (top) and validation of the transgene reporter (bottom) for **(A)** Insulin Red Nuclear (IRN) and **(B)** Amylin Nuclear Green in IRN (ANGIs) knock-in. **(C)** Transgene fluorescence of ANGIs reporter line at day 7 of stage 6 of SC-islet differentiation protocol quantified by flow cytometry. **(D)** Structured illumination image of ANGIs reporter line after SC-islet differentiation. (**E**) INS^mCherry^ expression during prolonged *in vitro* culture during at stage 6 of SC-islet differentiation quantified by flow cytometry. **(F)** Flow cytometry timecourse of IAPP^eGFP^ expression within the INS^mCherry^ population during prolonged *in vitro* culture during at stage 6 of SC-islet differentiation. **(G)** Reaggregates of ANGIs subpopulations after live cell sorting. **(H)** Hormone expression in INS^+^ IAPP^+^ vs INS^+^ IAPP^-^cells. n = 4 differentiations for INS^+^ IAPP^-^ and n = 3differentiations for INS^+^ IAPP^+^ **(I)** Comparison of the sorting strategies based on IAPP and CD49A.

INS^+^ IAPP^+^ and INS^+^ IAPP^-^ cells could be purified and reaggregated from live SC-islet cultures to form uniform clusters of a single cell type (**Figure 2G**). INS^+^ IAPP^+^ cells had higher expression of β cell hormones and lower expression of other endocrine hormones than INS^+^ IAPP^-^ cells (**Figure 2H**). Finally, we compared the sorting strategy based on INS and IAPP to the previously proposed sorting strategy for CD49A (ITGA1) (Veres *et al*., 2019). We found that INS^+^ CD49A^+^ cells were not enriched in IAPP^+^ cells. However, majority of INS^+^ IAPP^+^ SC-β cells also expressed CD49A (**Figure 2I)**. These observations were consistent with the previous studies suggesting that CD49A is a β cell surface marker but not a selective surface marker of mature SC-β cells expressing IAPP.

### Functional characterization of INS**^+^** IAPP**^+^** SC-**β** cells

INS^+^ IAPP^+^ cells have a smaller proportion of glucagon^+^ polyhormonal cells than INS^+^ IAPP^-^ cells (**Figure 3A**). The fraction of somatostatin^+^ and ghrelin^+^ polyhormonal cells was similarly reduced (data not shown). INS^+^ IAPP^+^ cells exhibited higher intracellular insulin content and secretion compared to INS^+^ IAPP^-^ cells (**Figure 3B and C**). In addition, the IAPP^eGFP+^ subpopulation of SC-β cells is more stable over time in the final stage of differentiation, as these cells retained INS^mCherry^ expression after 9 days of culture reaggregation (**Figure 3D**).

**Figure 3.**
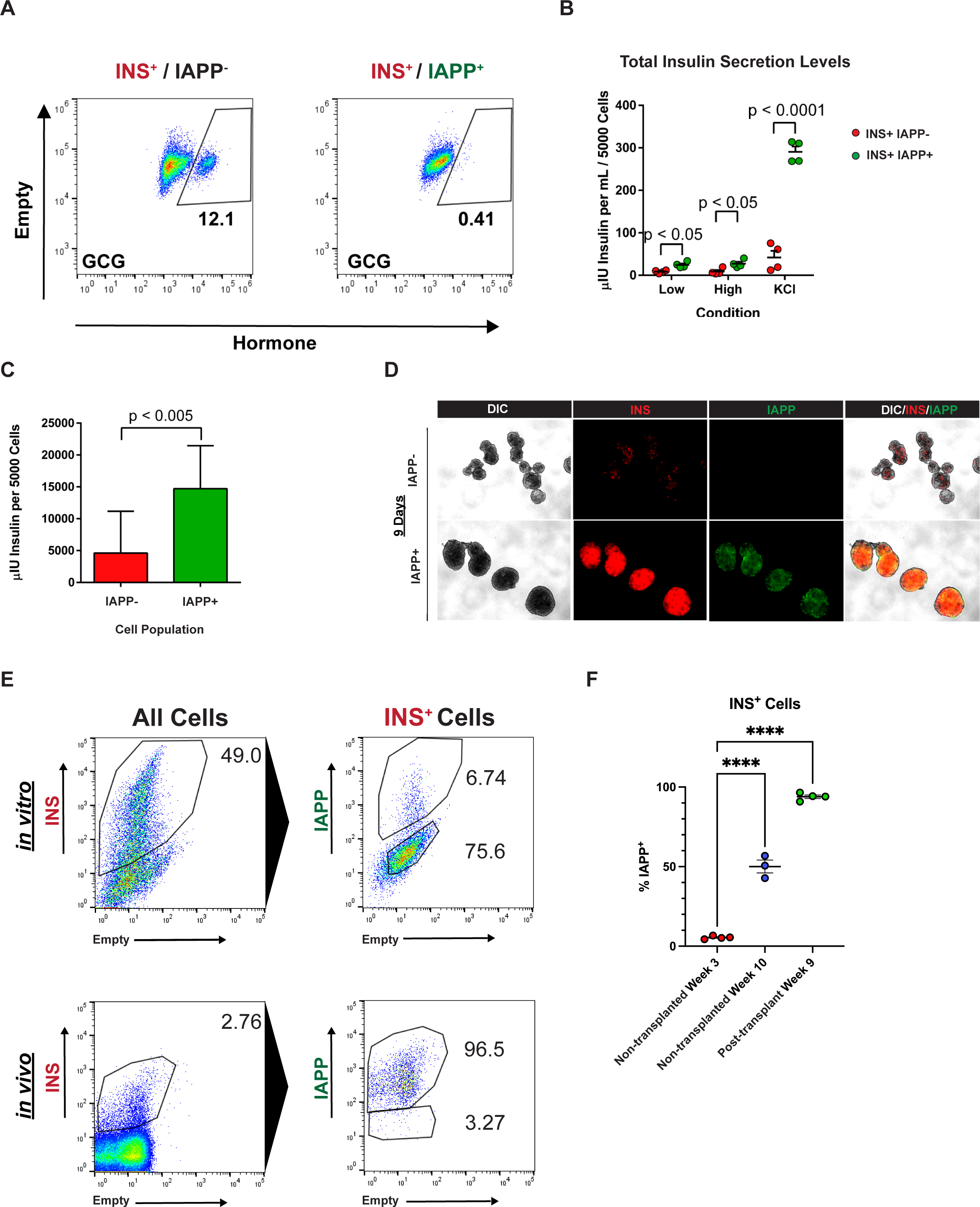
Characterization of INS^mCherry+^ IAPP^eGFP+^ vs INS^mCherry+^ IAPP^eGFP-^ cells. **(A)** Proportion of GCG^+^ bi-hormonal cells within INS^+^ population of SC-islets. **(B)** Insulin secretion under low glucose, high glucose and KCl-stimulated conditions. Statistical significance was determined using a 2-way ANOVA test. **(C)** Total insulin content per 5,000 cells in IAPP^eGFP+^ vs IAPP^eGFP-^ subpopulations in Stage 6 differentiated SC-islets. Significance was determined using a paired T-test. **(D)** Live cell epifluorescence for IAPP^eGFP+^ subpopulation reaggregates after 9 days of culture post-sorting. **(E)** Proportion of IAPP^eGFP+^ among INS^mCherry+^ cells *in vitro* and after 9 weeks *in vivo*. **(F)** Proportion of IAPP^eGFP+^ among INS^mCherry+^ cells in grafts harvested at 9 weeks *in vivo* and in cells maintained in culture in parallel. Significance was determined using a 2-way ANOVA test. (p < 0.0001). All experiments in panels A-D were carried out in SC-islets after two weeks of culture in stage 6 to allow for accumulation of INS^+^ IAPP^+^ cell population.

After profiling IAPP-expressing SC-β cells *in vitro*, we next sought to understand the dynamics of IAPP expression during maturation *in vivo*. We transplanted 5 million unsorted stage 6 dual reporter ANGIs SC-islets under the kidney capsule of NOD SCID immunocompromised mice and allowed the grafts to mature for 9 weeks. After transplantation, the grafts were dissociated and analyzed by flow cytometry. Cells from the same differentiations were kept in prolonged *in vitro* culture and analyzed in parallel. The fraction of IAPP^+^ cells among INS^+^ cells increased after transplantation *in vivo* (**Figure 3E**). Grafts harvested after transplantation exhibited an increase in IAPP^+^ INS^+^ cells among the INS^+^ population compared with the same differentiations maintained even longer *in vitro* (**Figure 3F**). Ultimately, nearly all INS^+^ cells gained IAPP expression at the end of 9 weeks *in vivo*.

### Aconitine induces IAPP Expression

We conducted a small molecule screen to identify compounds that increase the proportion of INS^+^ IAPP^+^ SC-β *in vitro*. Nine compounds increased IAPP expression in 2D culture (**Figure 4A, B**). Of these compounds, only aconitine was successfully validated in 3D culture and increased INS^+^ IAPP^+^ cells in a dose-dependent manner (**Figure 4C**). Up-regulation of IAPP was observed already within 48 hours of incubation with aconitine (**Figure 4D, E**). Addition of aconitine to the final stage of differentiation resulted in accelerated IAPP induction, with approximately double the number of IAPP^+^ cells after three weeks of culture (**Figure 4F**). Aconitine induced IAPP expression only in the INS^+^ cells, as no change in IAPP signal was observed in INS^-^ cells. Similarly, increased expression of IAPP was not accompanied by change in the overall INS^+^ population size (data not shown).

**Figure 4.**
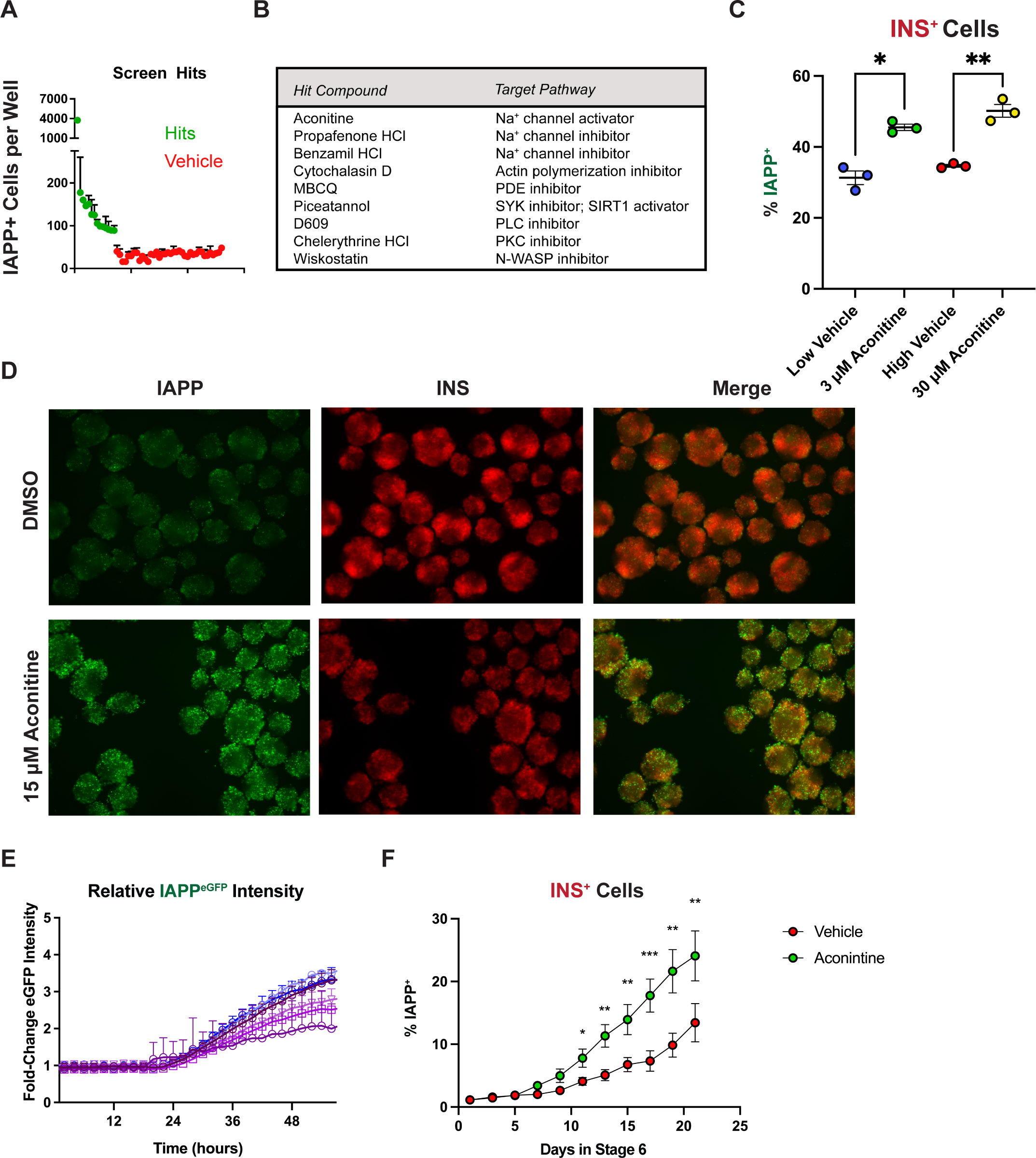
Aconitine induces IAPP expression. **(A)** IAPP^eGFP+^ nuclei counted per well in hits (green) and DMSO vehicle controls (red) after quantification of small molecule screen. **(B)** Identity and mechanism of action of the screening hits. **(C)** Dose response to aconitine in 3D culture after 96 hours of exposure to the compound. Low and high vehicle refer to matched DMSO concentrations for 3 and 30 µM aconitine conditions. Statistical significance was determined using a 1-way ANOVA test (* p < 0.05, ** p < 0.01.) (**D**) Epifluorescence image of live ANGIs clusters 48 hours after vehicle or 15 µM aconitine exposure. **(E)** Time course of IAPP^eGFP^ expression after short-term incubation with 15 µM aconitine, n = 6. Each line is the average eGFP intensity recorded across 30 wells normalized to DMSO control-treated wells. **(F)** Flow cytometry analysis of IAPP^eGFP+^ within INS^mCherry+^ cells with 15 µM aconitine or vehicle added to stage 6 SC-β differentiation protocol. Significance was determined using multiple Paired T-tests (* p < 0.01, ** p< 0.005.) Experiments in panels A-E were carried out in SC-islets cultured for 2 weeks in stage 6 to allow for accumulation of INS^+^ IAPP^+^ cells before exposure to aconitine.

## Discussion

In summary, the INS^+^ IAPP^+^ SC-β cell subpopulation was mono-hormonal, stable *in vitro* and *in vivo* and comprised the population of functional, insulin-producing cells after transplantation under the kidney capsule of immunocompromised mice. Thus, IAPP marks a more stable subpopulation of SC-β cells with higher insulin content that may be the source of insulin production after successful engraftment *in vivo*.

IAPP is often described as a diabetes-associated protein capable of forming amyloid fibrils that damage islets in type 2 diabetes (Westermark et al., 2011). However, IAPP is co-secreted with insulin in adult β cells (Westermark *et al*., 2011) and does not form fibrils in a heterodimer with insulin (Shen et al., 2006; Wiltzius et al., 2009). Thus, IAPP has a physiological and not only a pathological role and is a logical marker of maturing SC-β cells *in vitro*.

IAPP mRNA expressed in the pancreatic islet is β cell-specific and highly transcribed (Baron *et al*., 2016), which is beneficial for visualization using fluorescently encoded reporter transgenes, cell sorting, and screening purposes. We demonstrated feasibility of the approach by generating the dual reporter cell line ANGIs and conducting a small molecule screen. IAPP expression was induced in INS^+^ SC-β cells by incubation with aconitine at the final differentiation stage. Interestingly, extract from the aconite plant (not isolated aconitine) improves glycemic control and increases percentage of β cells on pancreatic histology slices in rats with streptozotocin-induced diabetes (Liou et al., 2006; Shoaib et al., 2020). By contrast, aconitine does not affect insulin secretion in INS1 and MIN6 cells at low and high glucose (Burns et al., 2015) and does not significantly increase percentage of PDX1^+^ progenitor cells in differentiating embryonic stem cells (Chen et al., 2009). Thus, our findings are in line with published literature indicating that aconitine increases the stability of β cells. Aconitine opens Na^+^ channels but also induces subsequent intracellular accumulation of calcium ions (Sun et al., 2014). Genes co-expressed with IAPP were calcium-dependent, supporting calcium accumulation as a potential mechanism of action.

IAPP has also recently been identified by other groups as a marker of more mature SC-β cells before and after transplantation (Augsornworawat et al., 2020; Bruin et al., 2015; Rezania *et al*., 2014; Rezania et al., 2012; Van Hulle et al., 2021). Our work supports these findings and expands on the utility of IAPP as a marker by functionally interrogating the IAPP-expressing subpopulation of SC-β cells. The connection between expression of a maturation-associated islet hormone and manipulation of membrane ion channels also suggests that manipulating membrane voltage and ion balances across the cell membrane may be a promising avenue to pursue further maturation of stem cell-derived islets *in vitro*. Further studies that improve IAPP induction *in vitro* and provide mechanistic insights into this desirable phenotype might contribute to generating an *in vitro* stem cell-derived therapy for diabetic patients.

## Methods

### Generation of 1016 iPS GFP Line

The 1016 iPS cell line was obtained from Columbia University. Constitutive GFP-expressing 1016 iPS cells were generated as described (Hockemeyer et al., 2011). Cells were selected using 2 µM puromycin, expanded, and adapted to 3D suspension culture. All experiments using human embryonic or other pluripotent stem cells were reviewed and approved by the Harvard University Embryonic Stem Cell Research Oversight (ESCRO) Committee.

### Generation of ANGIs cell line

1016 iPS cells were grown at low passage and electroporated using the Neon nucleofector system using 2, 1050 V pulses at 30 msec in the presence of px330 plasmid encoding Cas9 and containing guide RNAs targeting the stop codon of the insulin locus and a second donor vector containing homology 5’ and 3’ from the target site and encoding an in-frame P2A mCherry with antibiotic selection. Cells were allowed to recover for 72 hours followed by addition of antibiotic for positive selection of clones having undergone successful recombination. After approximately 10 days, large colonies were picked and grown in 96 well format, expanded, and tested using PCR primers targeting the insulin locus and the transgene. These insulin red nucleus (IRN) clones were then targeted using a second round of electroporation targeting the IAPP locus. Again, colonies were picked, PCR screened, and validated. Two individual clones were used for this manuscript, maintained as separate banks and individually differentiated for experiments.

### Donor islets

Islets were purchased from Prodo Laboratories, NDRI, and the University of Miami. All islets were isolated from healthy donors at the time of death and used within 7 days of shipment. All islet studies were performed in accordance with IRB approved procedures at Harvard University under protocol IRB16-0013.

### Directed differentiation of SC-**β** cells

Pluripotent iPS cells were cultured in suspension culture using mTeSR1 basal medium and passaged using accutase and differentiated using the protocol in (Pagliuca *et al*., 2014). Efficiency of SC-β cell induction was confirmed as in (Pagliuca *et al*., 2014). Stage 3 culture media was used for extended culture in stage 6.

### Immunofluorescence

Sections of SC-β clusters and cross-sections of pancreas were fixed in 4% paraformaldehyde for approximately 1 hour before washing in PBS and embedded in paraffin blocks for sectioning. Images were taken of 10 µm sections of tissue using a Zeiss Imager.Z2. Antibodies used for immunofluorescence were Rat Anti-C-peptide (Developmental Studies Hybridoma Bank, ID-4) and Goat Anti-GLP-2 (Santa Cruz Antibodies, C-20).

### Flow cytometry

Flow cytometry analysis of differentiation efficiencies was performed using the LSRII flow cytometer by BD. FACS purification of pre- and post-transplant SC-β cells was performed using a combination of MoFlo Astrios, MoFlo XDP, and BD FASCARIA III sorting instruments. Cells were first gated on side and forward scatter profiles followed by gating on GFP-positive cells and subsequent TSQ (405 nm)-positive PI-negative populations following the protocol (Davis *et al*., 2019). Cells were sorted into stage 6 medium and centrifuged/washed before lysis in TriZOL RNA reagent for extraction using trizol/chloroform extraction.

Flow cytometry for ANGIs differentiations were performed using the Attune NXT and sorted using the above-mentioned instruments. Gating strategies for ANGIs analysis was to first plot mCherry vs violet or empty channel to identify single positive cells and the mCherry^+^ subset was then plotted for eGFP vs empty channel or violet. Flow cytometry staining for ANGIs used the above antibodies as well as antibodies against endocrine hormones SST, PPY, GHRL in the far-red channel. These experiments included antibodies to mCherry and eGFP to enhance signal after fixation.

### Transplantation

The initial *in vivo* maturation experiments were carried out using immunocompromised SCID-Beige male mice from Jackson Laboratories (CB17.Cg-*Prkdc^scid^Lyst^bg-J^*/Crl). Surgeries were performed under the Melton protocol 10-18 for Harvard University. Animals were anesthetized using avertin and 5 million total cells of SC-β differentiations were transplanted underneath the left kidney capsule using a 23 gauge butterfly needle in ice cold RPMI 1640. Animals were allowed to recover on a heated pad under supervision until active before individual housing after surgery.

### Recovery of Transplanted Cells

Animals were sacrificed using CO_2_ asphyxiation and secondary cervical dislocation as per our animal protocol regulations. Individual kidneys were removed from the body cavity after sacrifice. Under a fluorescent dissecting microscope the kidney capsule was peeled from the body of the kidney, taking with it the GFP-positive graft (human cell reporter in the initial experiment) or INS^mCherry^-positive graft (experiments with the ANGIs cell line). After isolation in PBS, 1/3 of each graft was fixed in 4% PFA overnight and washed before embedding in paraffin wax and sectioned at a thickness of 10 μm for immunohistochemistry. The rest of the graft was incubated in accutase for 10 minutes at 37 degrees followed by passing through progressively higher gauge needles from 10/15/27 gauge. After dissociation cell suspensions were filtered through a 40 µM filter and resuspended in stage 6 medium and immediately sorted.

### In vitro GSIS

Pre-transplantation analysis of glucose stimulated insulin was performed using reaggregatres of 5000 cells in V-bottom ultra-low attachment 96 well plates. Clusters were fasted for 2 hours before analysis in 2.8 mM Glucose in Kreb’s ringer buffer (KRB) before addition of KRB at different glucose concentrations indicated or with 30 mM KCl in the presence of 2.8 mM glucose. Spent KRB was stored in 96 well plates frozen at −20 until analysis by insulin ELISA (Alpco Ultrasensitive Insulin ELISA) and read on a FluoSTAR Optima Spectrophotometer. Readings were taken at 450 and 650 nm and empty channel (650 nm) values were subtracted from 450 spectra to remove background absorbance.

### in vivo GSIS

After transplantation SCID-Beige mice were allowed to recover for 2 weeks followed by *in vivo* GSIS analysis at 2, 4 and 6 weeks post-transplant. Animals were fasted overnight prior to the GSIS assay. After overnight fast, blood was collected by a facial puncture, immediately followed by intraperitoneal injection of 2 g/kg glucose. After 30 minutes blood was again collected by the mandibular bleeding method. Blood was spun down and plasma isolated and frozen at −20 degrees Celsius until analysis by insulin ELISA.

### RNA Sequencing

Following RNA isolation, RNA quality was first assessed using Bioanalyzer RNA Pico chips. cDNA was then prepared using the Wafergen PrepX SPIA platform. Libraries were barcoded using Illumina indices and sequenced using the Illumina NextSEQ. Samples were sequenced using 75 bp paired-end reads. The reads were aligned to GRCh38 and quantified with RSEM (Li and Dewey, 2011). Read counts were TMM normalized (Robinson and Oshlack, 2010). Log_2_ (TMM+0.25) followed an approximately normal distribution based on visual examination of quantile-quantile plots against normal distribution. Differentially expressed genes post-vs pre-transplantation were identified using mixed effect linear model with cell differentiation batch as random intercept. Genes with Benjamini-Hochberg FDR < 0.05 were considered significant. GO term enrichment was analyzed online with PANTHER (http://pantherdb.org/) (Mi et al., 2017) and AmiGO2 (http://amigo.geneontology.org/amigo) (Carbon et al., 2009).

### Co-expression network

RNA-sequencing data on bulk human islets GSE50244 (Fadista et al., 2014) was reprocessed from FASTQ files with the bcbio pipeline (Guimera, 2011) with alignment to human genome build hg38 with hisat2 aligner and standard quality format. Counts were TPM-normalized and log2 transformed. MEGENA co-expression network (Song and Zhang, 2015) was constructed from genes with > 1 TPM in all samples.

### Small molecule screen

ANGIs cells were differentiated into week 1 of stage 6, followed by dispersion and re-plating on Matrigel in 384-well plates and incubated in compounds for 96 hours in basal stage 6 medium followed by high content imaging analysis. Each compound was included in two concentrations with four replicates per condition. Small molecule libraries were SCREEN-WELL compound libraries for nuclear receptor ligands, bioactive lipids, and Kinase inhibitors from Enzo Life Sciences. Compounds were fed twice, 48 hours apart and imaged using the Zeiss Cell Discoverer and downstream analyses were performed using the Zeiss Blue image analysis software. Aconitine-mediated upregulation of IAPP positivity was validated over short-term time course using IncuCyte^®^ live cell imaging to quantify the proportions of IAPP^eGFP^ reporter-positive cells among the INS^mCherry^ reporter-positive population.

## Data availability

RNA-seq experiments: GSE135944 of TSQ^+^ PI^-^ sorted SC-islet cells before and after *in vivo* maturation as well as human cadaveric islets and GSE263366 of INS^mCherry+^, IAPP^eGFP+/-^ SC-β cells after 14 days of culture in stage 6. The co-expression network model is included as supplementary data S1 and can be read into R with the igraph package.

## Author contribution statement

Project management, resources and funding acquisition: B.T., D.A.M. *in vitro* and *in vivo* experiments: J.C.D., J.H.K., P.Ö.E. Computational analyses: J.C.D., M.R., S.M.. First draft: J.C.D. Review, editing, approval of the final version: all authors.

## Competing interest

M.R. and B.T. are current and S.M. were former employees of AstraZeneca when this research was conducted. AstraZeneca provided salaries for these authors, but had no role in conceptualization of the study, data collection, interpretation or preparation of the manuscript. The authors receive no financial or non-financial reward for publication. D.A.M. is the founder of Semma Therapeutics and is now an employee of Vertex Pharmaceuticals, which has licensed technologies from Harvard and HHMI. J.H.K. is now a Vertex employee. All work was conducted at Harvard University with research cell lines. Other authors declare no competing interests. A patent related to this work was filed by Harvard University.

## Acknowledgements

This work was supported by grants from the National Institutes of Health and the National Institute of Digestive and Diabetic Kidney Disease Human Islet Research Network UC4 DK104159. We thank Edwin Rosado-Olivieri and Aharon Helman for experimental support and additionally Stephanie Tsai, Jose Rivera-Feliciano, Nils Bergenhem, Nadav Sharon, Ramona Pop, Mårten Hammar and Can Kayatekin for helpful discussion. All of these experiments were done at Harvard University and we would like to acknowledge The Bauer Core Facility at Harvard University, Jeffrey Nelson and Zachary Niziolek, the Harvard Center for Biological Imaging (RRID:SCR_018673), Douglas Scott Richardson, Christian Hellriegel, and the Biopolymers Facility at Harvard Medical School for logistics and support.

## Notes

https://www.ncbi.nlm.nih.gov/geo/query/acc.cgi?acc=GSE135944

https://www.ncbi.nlm.nih.gov/geo/query/acc.cgi?acc=GSE263366

